# Evaluating the efficacy of prototype antiseizure drugs using a preclinical pharmacokinetic approach

**DOI:** 10.1101/2022.07.07.499055

**Authors:** Jeffrey A. Mensah, Kristina Johnson, Christopher A. Reilly, Karen S. Wilcox, Joseph E. Rower, Cameron S. Metcalf

**Affiliations:** Epilepsy Therapy Screening Program (ETSP) Contract Site, University of Utah, Salt Lake, UT, USA; Department of Pharmacology & Toxicology, University of Utah, Salt Lake City, UT, USA; Center for Human Toxicology, College of Pharmacy, University of Utah, Salt Lake City, UT, USA

**Keywords:** epilepsy, antiseizure drugs, preclinical rodent models, pharmacokinetics, efficacy

## Abstract

**Objective:** Pharmacokinetics (PK) of a drug drive its exposure, efficacy, and tolerability. A thorough preclinical PK assessment of antiseizure medications (ASMs) is therefore essential to evaluate the clinical potential. We tested protection against evoked seizures of prototype ASMs in conjunction with analysis of plasma and brain PK as a proof-of-principle study to enhance our understanding of drug efficacy and duration of action using rodent seizure models.

**Methods:** *In vivo* seizure protection assays were performed in adult male CF-1 mice and Sprague-Dawley rats. Clobazam (CLB), N-desmethylclobzam (NCLB), carbamazepine (CBZ), carbamazepine-10,11-epoxide (CBZE), valproic acid (VPA), and levetiracetam (LEV) concentrations were quantified in plasma and brain using liquid chromatography-tandem mass spectrometry. Mean concentrations of each analyte were calculated and used to determine PK parameters via non-compartmental analysis in Phoenix WinNonLin.

**Results:** NCLB concentrations were approximately 10-fold greater than CLB in mice. The antiseizure profile of CLB was partially sustained by NCLB in mice. CLB concentrations were lower in rats than in mice. CBZE plasma exposures were approximately 70% of CBZ in both mice and rats, likely contributing to the antiseizure effect of CBZ. VPA showed a relatively short half-life in both mice and rats, which correlated with a sharp decline in efficacy. LEV had a prolonged brain and plasma half-life, associated with a prolonged duration of action in mice.

**Significance:** The study demonstrates the utility of PK analyses for understanding the seizure protection time-course in mice and rats. The data indicate that distinct PK profiles of ASMs between mice and rats likely drive differences in drug efficacy between rodent models.

**Key Points:**

- There exist potential contributions of active metabolites to the efficacy of some ASMs.
- The utility of preclinical PK assessment of ASM is critical to guide our insight into a drug efficacy profile and provide a framework for subchronic dosing strategies.
- Species-specific variations in PK profiles of ASMs in rodent models of epilepsy may underpin the differences in antiseizure effect in these models.
- Pre-clinical drug screening of ASMs should include a (sub)chronic dosing paradigm to better mimic the dosing regimen in the clinic.

## Introduction

Antiseizure medications (ASMs) are the most common therapeutic approach for treating seizures in epilepsy. However, pharmacotherapy shortcomings in seizure control are well known. Approximately 30% of patients with epilepsy are pharmacoresistant to current ASMs.^1^ ASMs also carry dose-limiting side effects that can significantly hinder their chronic use by patients.^2^ Due to the limitations associated with existing ASMs; there is a need for continued research into discovering novel ASMs with improved efficacy and safety profiles.

Typical preclinical screening and evaluation methods for novel ASMs utilize primarily mouse and/or rat models to assess compound safety and protection against seizures.^3,4^ Frequently, the onset, duration, magnitude of drug efficacy, and tolerability of ASMs, vary between various rodent seizure models.^5-8^ One possible explanation for the observed variability in preclinical outcomes is that the pharmacokinetic (PK) properties of ASMs differ between species and strains of rodents. In fact, previous studies demonstrated significant intra-and interspecies variation in PK profiles of common ASMs, including clobazam (CLB), valproic acid (VPA), carbamazepine (CBZ), and levetiracetam (LEV).^5,9,10^ Additionally, poor preclinical understanding of a compound’s PK may be an underlying reason for difficulties translating ASMs with preclinical success in the clinic setting. Therefore, preclinical PK-PD studies will provide vital information for subsequent drug development and the clinical utility of ASMs.^11^

The Epilepsy Therapy Screening Program (ETSP) of the National Institute of Neurological Disorders and Stroke (NINDS) utilizes well-established rodent epilepsy models to identify and profile novel compounds as potential new treatments for drug-resistant epilepsy. Rational evaluation of the preclinical efficacy of ASMs must include an understanding of plasma and brain PK. Evaluating brain and plasma PK to understand safety and efficacy outcomes across mouse and rat seizure models is an integral part of early drug discovery or development work and a priority of the ETSP. The absence of these data limits the translation of preclinical findings to the clinic and delays identifying an optimal human dose of a novel ASM. This study begins to fill this fundamental gap in our knowledge by demonstrating the utility of evaluating plasma and brain PK profiles to guide efficacy studies using rodent seizure models. Moreover, data from the study will guide informed decisions about subchronic dosing strategies that will provide an excellent framework for the ETSP to begin subchronic administration and screening of new investigational compounds, starting with prototype ASMs; CLB, VPA, CBZ, and LEV, which constitute four important reference compounds in the ETSP’s drug-resistant testing workflow.

## Materials and Methods

### Animals

All animal care and experimental procedures were approved by the Institutional Animal Care and Use Committee of the University of Utah. Adult male CF No 1 (CF-1) albino mice (18 - 30 g) and adult male Sprague-Dawley rats (100 -150 g) were obtained from Charles River’s facility in Kingston, NY, U.S.A. The animals were housed, fed, and handled in a manner consistent with the recommendations in the National Council Publication, “Guide for the Care and Use of Laboratory Animals.” All mice and rats were housed in plastic cages in specially constructed rooms with controlled humidity, exchange of air, and controlled lighting (12 hrs on - 12 hrs off). Animals newly received in the laboratory are allowed sufficient time to correct for possible food and water restriction incurred during transit before being employed in testing.

### Drug dose selection, preparation, and administration

CBZ, CLB, and VPA were purchased from Sigma-Aldrich, Inc. (St. Louis, MO). LEV was obtained from T.C.I. America (Portland, OR). All drugs used in this study were formulated using 0.5% methylcellulose (Sigma, St. Louis, MO) suspensions and administered using dose volumes of 10 mL/kg (mice) or 4 mL/kg (rats). Generally, we selected drug doses known to be effective (ED_50_) in the seizure assays and a second dose near the upper limit of tolerability tolerated (TD_50_) to provide a dose range that can produce a reasonable therapeutic concentration range and inform doses for subchronic administration. The ED_50_ for each drug was determined from previous studies using the Probit analysis^12,^ whereas TD_50_ was determined using the rotarod test for mice and the minimum motor impairment test for rats^13^ in our lab. All compounds were single-dose via intraperitoneal (i.p.) injections. The dose and the rodent seizure assay used to evaluate the efficacy of each drug are listed in **Table 1**.^5,11,13-15^

**Table 1:** List of prototype ASDs and selected doses (ED_50_ -low dose and TD_50_ -high dose) evaluated in mouse and rat seizure test models.

|  |  |  |  |
| --- | --- | --- | --- |
| Compound | Species | Dose (mg/kg) *a,b* | Efficacy test |
| CLB | Mouse | 7 | 6 Hz 44 mA |
|  |  | 17 |  |
|  | Rat | 17 | MES |
|  |  | 37 |  |
| CBZ | Mouse | 7 | MES |
|  |  | 40 |  |
|  | Rat | 3 |  |
|  |  | 24 |  |
| VPA | Mouse | 260 | 6 Hz 44 mA |
|  |  | 570 |  |
|  | Rat | 165 | MES |
|  |  | 315 |  |
| LEV | Mouse | 20 | 6 Hz 44 mA |
|  |  | 500 |  |
|  | Rat | 14 | 6 Hz 80 V |
|  |  | 800 |  |

CLB-Clobazam; CBZ-Carbamazepine; VPA-Valproic acid; LEV-Levetiracetam *a,b* Values from Ref (6, 11, 13-15)

### Seizure Test Models

#### Maximal Electroshock test (MES)

Compounds (CBZ, VPA, and CLB) were tested for their antiseizure effects using the MES assay following i.p. administration to adult CF-1 mice and adult Sprague-Dawley rats. Tetracaine (2-3 drops) was applied to the eyes of each animal before stimulation. 60 Hz of alternating current (50 mA in mice and 150 mA in rats) was delivered for 0.2 seconds by corneal electrodes at various selected time points after drug treatment. An animal was considered “protected” from MES-induced seizures upon abolishing the hindlimb tonic extensor component of the seizure.

#### 6 Hz 44 mA test

We assessed the efficacy of compounds (CLB, VPA, and LEV) in the 6 Hz 44 mA assay in adult male CF-1 mice. Briefly, 2-3 drops of tetracaine were applied to animals’ eyes. A 44 mA current (two times the convulsive current producing seizures in 97% of mice CC_97_), as previously described by Metcalf et al.^13^ and Barton et al.^5^ was then delivered for 3 seconds by corneal electrodes in drug pretreated mice at various selected time-points. An initial momentary stun characterized seizures followed immediately by jaw clonus, head nodding, forelimb clonus, twitching of the vibrissae, and Straub tail lasting for at least 1 second.

#### 6 Hz 80 V test

LEV efficacy against pharmacoresistant seizures in male adult Sprague-Dawley rats was tested using a 6 Hz 80 V assay. 2-3 drops of tetracaine were applied to the eyes of animals. Delivery of 6 Hz, 0.2 msec rectangular pulse, 3 seconds duration by a corneal electrode (80 V) was performed following pretreatment with test compound at various selected time-points. Seizures were characterized and recorded as head nodding, jaw clonus, forelimb clonus with occasional bilateral forelimb clonus, rearing, and loss of the righting reflex.

#### Blood and brain sample collection

Blood samples were collected from animals (4 animals per time point) immediately after conducting behavioral assays at 0.25, 0.5, 1, 2, 4, and 8 hr post-dose. Trunk blood was collected via decapitation into tubes with K_2_EDTA (Becton, Dickson and Company, NJ) as an anticoagulant. Plasma was isolated from blood using the accuSpin Micro 17R centrifuge (3500 x g for 10 min at 4°C). Brains were rapidly removed following decapitation from the same animal and weighed. Whole brains were homogenized in ultrapure water using a probe sonicator to obtain a target homogenate concentration of 200 mg tissue/mL. Plasma and brain homogenate samples were stored at −80°C until analysis.

#### Bioanalysis

CBZ, CBZ-10,11-epoxide (CBZE), CLB, N-desmethyl clobazam (NCLB), VPA, LEV, CBZ-^13^C_6_, CBZE-^13^C_6_, CLB-^13^C_6_, NCLB-^13^C_6_, VPA-^13^C_6_, and LEV-^13^C_6_ were purchased from Sigma-Aldrich, Inc. (St. Louis, MO). NCLB, NCLB-^13^C_6_, LEV, LEV-^13^C_6_, and VPA-^13^C_6_ were purchased as pre-diluted 1 mg/mL stock solutions. All others were purchased in powder form and diluted with 50:50 methanol: water (MeOH:H_2_O) to 1 mg/mL, except for VPA, which was prepared at 10 mg/mL. Both plasma and brain were extracted and analyzed using the same assay. Plasma samples were diluted 1:10 (i.e., 10 µL sample with 90 µL of analyte-free plasma, total volume 100 µL) before analysis; a 100 µL aliquot of brain homogenate was extracted without dilution.

Sample extraction used a basified liquid-liquid procedure in borosilicate tubes. An internal standard solution (in 50:50 MeOH:H2O) containing isotopically labeled analogs of each analyte was added to every sample after aliquoting (CBZ-^13^C_6_, CBZE-^13^C_6_ and LEV-^13^C_6_ at 0.1 µg/mL; CLB-^13^C_6_ and NCLB-^13^C_6_ at 0.01 µg/mL; VPA-^13^C_6_ at 0.5 µg/mL final concentrations). The sample was then basified using 0.5 mL of 5 mM ammonium bicarbonate (NH_4_HCO_3_, pH 8) solution and extracted using 2.0 mL of methyl tert-butyl ether (MTBE). The sample was then vortex mixed, centrifuged, and frozen before decanting and evaporating the MTBE layer under house air (15 psi) in a TurboVap set at 40°C. The sample was then reconstituted with 200 µL of 5:95 MeOH:H2O and transferred into an autosampler vial for instrumental analysis.

Analysis of CBZ, CBZE, CLB, and NCLB utilized a ThermoScientific TSQ Vantage tandem mass spectrometer (MS/MS) coupled with a ThermoScientific Accela liquid chromatography (LC) and autosampler system. VPA and LEV analyses were conducted on a ThermoScientific TSQ Quantum Access MS/MS coupled with an Agilent 1100 LC and autosampler. Analyte separation on both instruments was achieved using gradient elution with mobile phases consisting of (A) 5 mM NH_4_HCO_3_, pH=8 and (B) MeOH on a Phenomenex Luna Omega PS C18, 3 µm (100 × 2.1 mm) column. Quantitation used the following mass transitions (collision energy, CE): 237.2→194.1 (CBZ; CE=26 V); 243.2→200.1 (CBZ-^13^C_6_, CE=19 V); 253.1→180.1 (CBZE, CE=24 V); 259.1→186.1 (CBZE-^13^C_6_, CE=34 V); 287.1→245.1 (NCLB, CE=19 V); 293.1→251.1 (NCLB-^13^C_6_, CE=19 V); 301.1→259.1 (CLB, CE=25 V); 307.1→265.1 (CLB-^13^C_6_, CE=19 V); 171.1→126.1 (LEV, CE=15 V); 177.1→132.1 (LEV-^13^C_6_, CE=15 V); 143.2→143.2 (VPA, CE=20 V); and 149.2→149.2 (VPA-^13^C_6_, CE=20 V). The dynamic ranges of the assays were 0.02 to 4.0 µg/mL (CBZ, CBZE, LEV), 2.0 to 400 ng/mL (CLB, NCLB), and 0.5 to 20 µg/mL (VPA).

#### PK Analysis

Before PK analyses, concentrations were averaged for each species, dose, and tissue combination at each time point. PK analyses were conducted in Phoenix WinNonLin v8.3, using the linear trapezoidal linear interpolation option calculation method in the non-compartmental analysis (NCA) module. The software was allowed to determine the best fit line for the elimination slope for all calculations. The parameters of interest for this analysis were the area under the concentration-time curve from 0 to 8 hr (AUC_0-8hr_), maximum concentration (C_max_), minimum concentration (C_min_), time to reach maximum concentration (T_max_), and half-life (t_1/2_). AUC_0-8hr_ was used to calculate metabolite: drug ratios for CBZ and CLB, while AUC_0-8hr_ and C_max_ were used to assess dose proportionality.

## Results

### CLB

The plasma t_1/2_ of CLB in mice was short and similar between the low (t_1/2p_ = 0.704 hr) and high (t_1/2p_ = 0.793 hr) dose cohorts. In contrast, NCLB exhibited a moderately long t_1/2_ in mouse plasma (low-dose t_1/2p_ = 6.41 hr and high-dose t_1/2p_ = 4.43 hr), resulting in NCLB having a plasma area under the curve (AUC) approximately 10-fold greater than CLB. Similarly, NCLB exhibited a long t_1/2_ in the mouse brain **(Figure 1A)**. Efficacy data from the 6Hz 44 mA test in mice are shown in **Table 2**. In the low-dose CLB mouse arm, the loss of efficacy mirrored the decline in CLB concentrations in the brain **(Table 2 and Figure 1A)**. As expected, greater effectiveness was observed in the high-dose CLB-treated mice, likely partially sustained by the high NCLB brain concentrations **(Table 2 and Figure 1A)**.

**Table 2:** Efficacy of CLB, VPA, LEV in the 6 Hz 44 mA and CBZ in the MES models of seizure test following intraperitoneal administration to adult male CF-1 mice.

|  |  |  |  |  |  |  |  |
| --- | --- | --- | --- | --- | --- | --- | --- |
| Compound | Dose (mg/kg) | Efficacy in mice (number seizure protected/number tested) |  |  |  |  |  |
|  |  | Time test (h) |  |  |  |  |  |
|  |  | 0.25 | 0.5 | 1 | 2 | 4 | 8 |
| CLB | 7 | 2/4 | 1/4 | 3/4 | 0/4 | 0/4 | 0/4 |
|  | 17 | 4/4 | 4/4 | 3/4 | 3/4 | 1/4 | 0/4 |
| CBZ | 7 | 3/4 | 3/4 | 2/4 | 0/4 | 0/4 | 0/4 |
|  | 40 | 4/4 | 4/4 | 4/4 | 4/4 | 4/4 | 0/4 |
| VPA | 260 | 4/4 | 3/4 | 0/4 | 0/4 | 0/4 | 0/4 |
|  | 570 | 4/4 | 4/4 | 4/4 | 4/4 | 0/4 | 0/4 |
| LEV | 20 | 0/4 | 0/4 | 0/4 | 0/4 | 0/4 | 0/4 |
|  | 500 | 1/4 | 1/4 | 1/4 | 1/4 | 1/4 | 0/4 |

**Figure 1:**
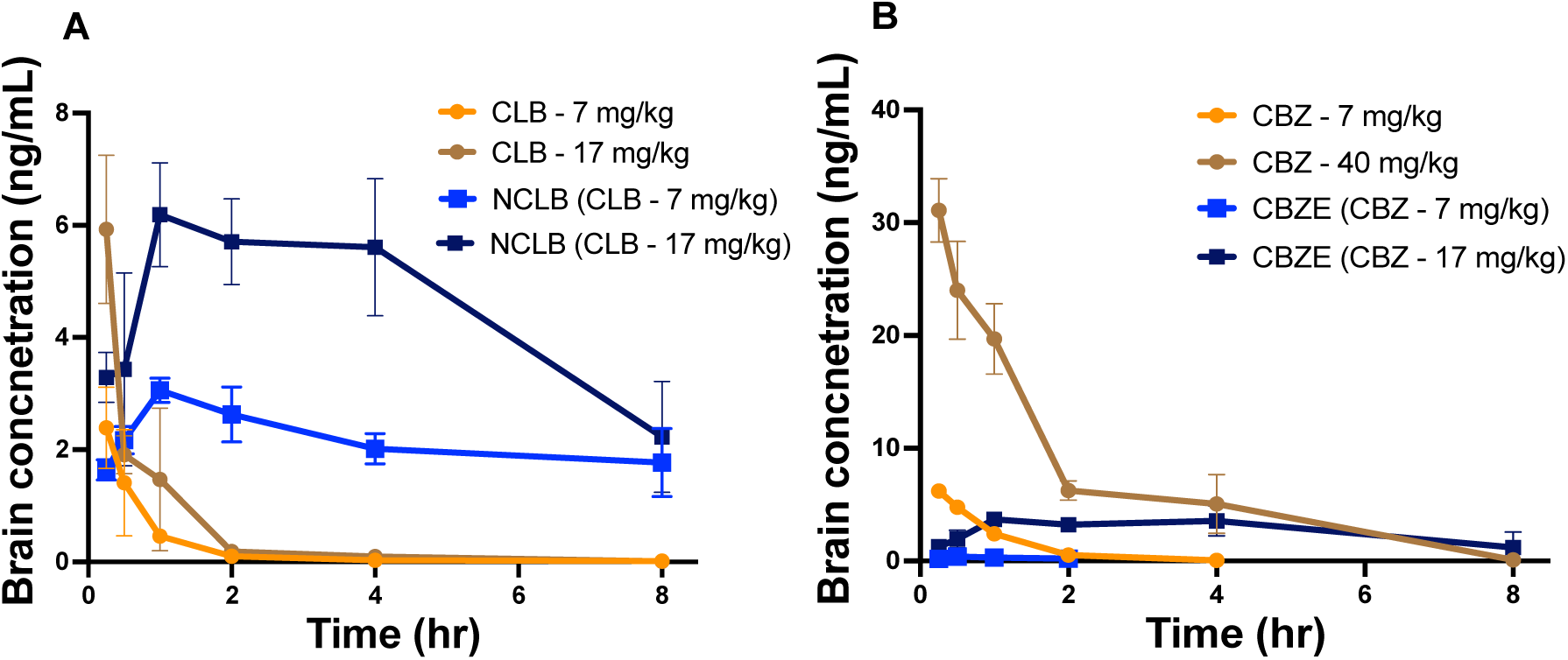
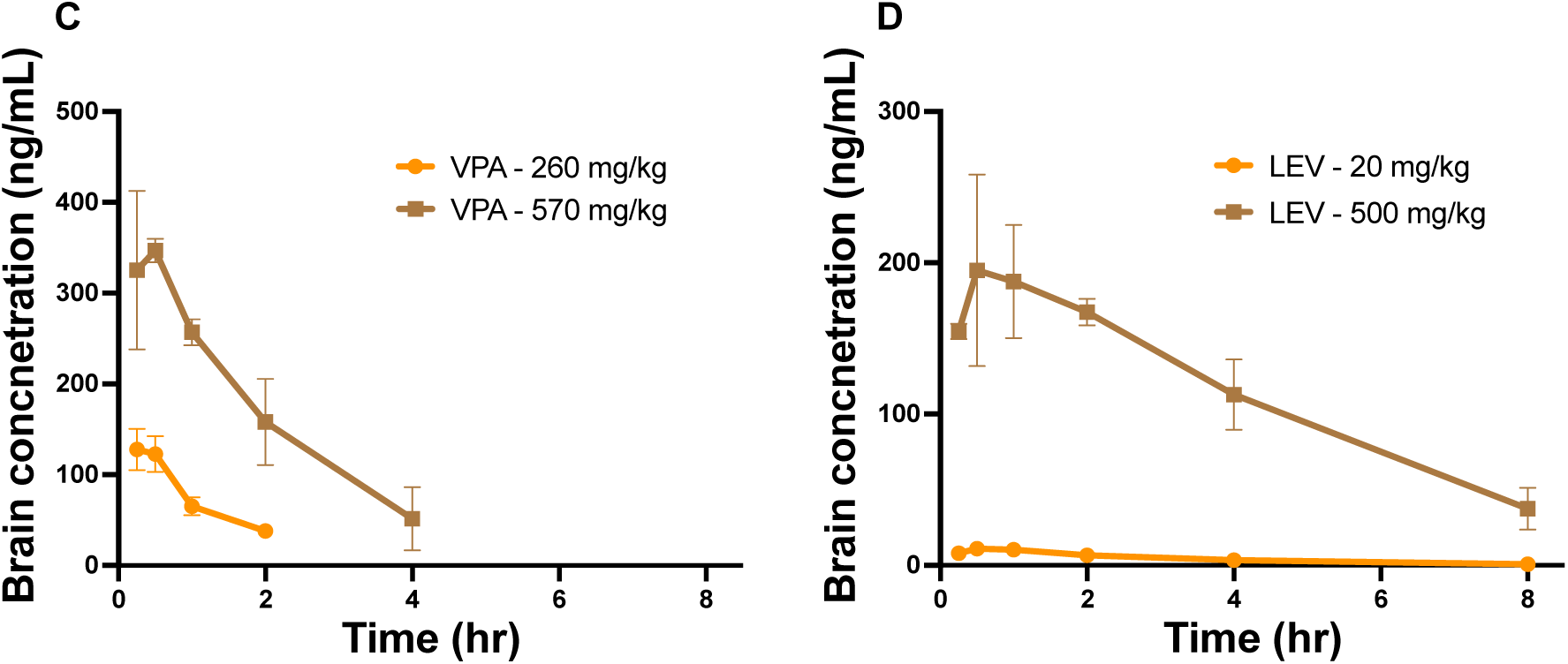
Concentration-time curves of (A) CLB and NCLB, (B) CBZ and CBZE, (C) VPA, and (D) LEV in the brain of adult male CF-1 mice following intraperitoneal administration. Brain samples were obtained between 0-8 hr post-dose. Each data point is the mean brain concentration value for 4 mice.

In rats, the t_1/2_ of CLB and NCLB was short in both plasma and brain tissue, with NCLB particularly short-lived in rat plasma.. In contrast to data from mice, NCLB concentrations were substantially lower (∼20 fold less) than CLB in plasma and brain in rats, indicative of differential metabolism rates between the two species. Similar to the mouse data, the kinetic profiles of CLB and NCLB in the rat plasma and brain were identical. The low-dose CLB did not protect rats from seizures at any time post-dose using the MES test **(Table 3)**. In the rats treated with a high-dose CLB, seizure protection was observed within the first hour after treatment but was absent after that **(Table 3)**, consistent with the rapid elimination rate of CLB from the brain in rats.

**Table 3:**
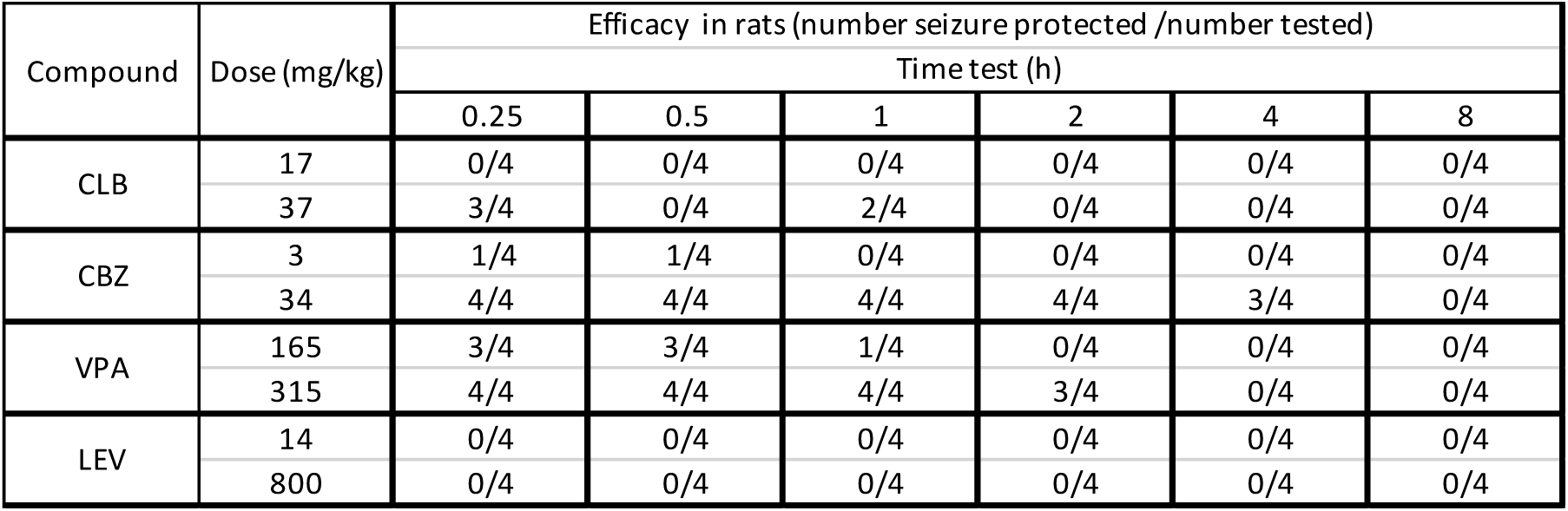
Efficacy of CLB, CBZ, VPA in the MES and LEV in the 6 Hz 80 V models of seizure test following intraperitoneal administration to adult male sprague dawley rats.

### CBZ

CBZ plasma half-life in the low-dose (t_1/2p_ = 1.30 hr) mice was short and similar to that of the high-dose (t_1/2p_ = 1.11 hr). CBZE was determined to have a slightly longer t_1/2_ in mouse plasma for the high-dose (t_1/2p_ = 3.98 hr) cohort relative to the low-dose (t_1/2p_ = 1.20 hr) arm. In mice, high-dose CBZE exhibited a longer plasma half-life (∼3-fold longer) than CBZ. CBZE concentrations were lower than CBZ in all matrices/doses. Additionally, high-dose CBZ exposures were approximately 50 - 60% greater than expected from low-dose, suggesting potential saturable clearance. A reduction in drug efficacy correlated with a decrease in CBZ mouse brain concentrations in the low-dose CBZ **(Figure 1B)**. In high-dose CBZ-treated mice, complete protection against seizure activities was observed to be rapid and sustained over the majority of the time course **(Figure 1B)**. The prolonged CBZ protective effect in the high-dose arm was observed despite the rapid elimination of parent CBZ, indicating that sustained CBZE exposure may add to the efficacy of CBZ.

The t_1/2_ of CBZ and CBZE in rats was generally short, though CBZE was found to have half-lives of approximately 4 hrs in both plasma and brain **(Table 4)**. High-dose CBZ concentrations were approximately two-fold greater than expected from low-dose, suggesting that CBZ was not dose-proportional in rats. Low-dose CBZ efficacy was minimal (25% protected), observed only within the first hour after drug treatment in the MES test **(Table 3)**. 100% seizure protection was achieved rapidly in the high-dose rat cohort and was sustained for a longer period (∼2 hrs) relative to the low-dose rat group, consistent with the higher brain concentrations achieved in this group.

**Table 4:** Pharmacokinetic parameters derived following intraperitoneal administration of CLB, CBZ, VPA and LEV to adult male CF-1 mice and sprague dawley rats.

|  |  |  |  |  |  |  |  |  |  |
| --- | --- | --- | --- | --- | --- | --- | --- | --- | --- |
| Species |  | Mouse |  |  |  |  |  |  |  |
| Compound |  | CLB |  | CBZ |  | VPA |  | LEV |  |
| Dose (mg/kg) |  | 7 | 17 | 7 | 40 | 260 | 570 | 20 | 500 |
| Matrix | PK Parameters | Data are mean values |  |  |  |  |  |  |  |
| Plasma | AUC (ng*hr/mL) | 973 | 2263 | 3.36 | 28.6 | 746 | 2422 | 33.6 | 1064 |
|  | Drug:Metab | 9.32 | 9.84 | 0.29 | 0.76 | n/a | n/a | n/a | n/a |
|  | Cmax (ng/mL) | 1459 | 3786 | 3.15 | 16.50 | 417 | 789 | 15.9 | 397 |
|  | Cmin (ng/mL) | blq | 61.1 | 0.039 | 0.093 | 13.9 | 22.7 | 0.495 | 25.5 |
|  | Tmax (hr) | 0.25 | 0.25 | 0.25 | 0.25 | 0.25 | 0.50 | 0.25 | 0.25 |
|  | t1/2 (hr) | 0.704 | 0.793 | 1.30 | 1.11 | 0.732 | 1.33 | 1.72 | 2.05 |
|  | Dose Prop (AUC) | n/a | 0.958 | n/a | 1.49 | n/a | 1.48 | n/a | 1.27 |
|  | Dose Prop (Cmax) | n/a | 1.068 | n/a | 0.917 | n/a | 0.863 | n/a | 1.00 |
| Brain | AUC (ng*hr/mg) | 1.74 | 3.89 | 6.01 | 56.4 | 146 | 692 | 35.5 | 917 |
|  | Drug:Metab | 9.38 | 9.41 | 0.195 | 0.387 | n/a | n/a | n/a | n/a |
|  | Cmax (ng/mg) | 2.26 | 5.93 | 6.20 | 31.1 | 128 | 347 | 11.0 | 195 |
|  | Cmin (ng/mg) | 0.014 | 0.012 | 0.045 | 0.108 | 37.9 | 51.3 | 0.683 | 37.4 |
|  | Tmax (hr) | 0.25 | 0.25 | 0.25 | 0.25 | 0.25 | 0.50 | 0.50 | 0.50 |
|  | t1/2 (hr) | 2.26 | 1.51 | 0.52 | 0.99 | 0.926 | 1.28 | 1.81 | 2.73 |
|  | Dose Prop (AUC) | n/a | 0.92 | n/a | 1.64 | n/a | 2.17 | n/a | 1.03 |
|  | Dose Prop (Cmax) | n/a | 1.08 | n/a | 0.88 | n/a | 1.24 | n/a | 0.71 |
| Species |  | Rat |  |  |  |  |  |  |  |
| Compound |  | CLB |  | CBZ |  | VPA |  | LEV |  |
| Dose (mg/kg) |  | 17 | 37 | 3 | 24 | 165 | 315 | 14 | 800 |
| Matrix | PK Parameters | Data are mean values |  |  |  |  |  |  |  |
| Plasma | AUC (ng*hr/mL) | 1453 | 4059 | 2.03 | 37.2 | 237 | 713 | 71.8 | 2274 |
|  | Drug:Metab | 0.036 | 0.048 | 0.695 | 0.672 | n/a | n/a | n/a | n/a |
|  | Cmax (ng/mL) | 1242 | 2829 | 2.13 | 19.4 | 241 | 415 | 18.6 | 529 |
|  | Cmin (ng/mL) | blq | blq | blq | 0.194 | 17.8 | 11.2 | 4.72 | 97.4 |
|  | Tmax (hr) | 0.25 | 0.25 | 0.25 | 0.25 | 0.25 | 0.50 | 0.50 | 0.50 |
|  | t1/2 (hr) | 2.60 | 1.89 | 0.82 | 1.18 | 0.417 | 0.616 | 6.16 | 3.19 |
|  | Dose Prop (AUC) | n/a | 1.28 | n/a | 2.29 | n/a | 1.58 | n/a | 0.55 |
|  | Dose Prop (Cmax) | n/a | 1.047 | n/a | 1.138 | n/a | 0.902 | n/a | 0.500 |
| Brain | AUC (ng*hr/mg) | 2.88 | 7.86 | 3.50 | 52.8 | 52.0 | 183 | 53.4 | 1719 |
|  | Drug:Metab | 0.045 | 0.054 | 0.420 | 0.481 | n/a | n/a | n/a | n/a |
|  | Cmax (ng/mg) | 2.90 | 6.59 | 2.91 | 28.3 | 78.6 | 145 | 8.48 | 307 |
|  | Cmin (ng/mg) | 0.027 | 0.051 | 0.088 | 0.322 | 27.0 | 43.4 | 3.52 | 90.6 |
|  | Tmax (hr) | 0.25 | 0.25 | 0.25 | 0.25 | 0.25 | 0.25 | 0.50 | 1.00 |
|  | t1/2 (hr) | 1.54 | 1.25 | 0.77 | 1.20 | 0.467 | 0.895 | 4.42 | 3.53 |
|  | Dose Prop (AUC) | n/a | 1.25 | n/a | 1.89 | n/a | 1.84 | n/a | 0.56 |
|  | Dose Prop (Cmax) | n/a | 1.044 | n/a | 1.216 | n/a | 0.968 | n/a | 0.630 |

### VPA

**Table 4** summarizes the PK parameters of VPA in plasma and brain for both mice and rats. The plasma half-life of VPA in mice was short and similar between the low-(t_1/2p_ = 0.73 hr) and high-dose (t_1/2p_ = 1.33 hr) VPA-treated mice. Similar half-lives were found in plasma and brain tissue. VPA efficacy data in mice from the 6 Hz 44 mA test are described in **Table 2**. In the low-dose VPA mouse arm, seizure protection was observed within the first hour of drug treatment before a rapid loss of efficacy mirroring the decline in VPA brain concentrations in mice. The high-dose VPA-treated mice showed maximal antiseizure efficacy, sustained only within 2 hrs, but absent after that. Data comparison from the two-dose levels suggests a need for brain concentrations of VPA to remain above 100 ng/mg for complete seizure protection.

The t_1/2_ of VPA in the plasma and brain of rats was extremely short (<1 hr), with total drug elimination (i.e.,>97% drug eliminated) occurring in less than 4.5 hrs **(Table 4)**. Efficacy results from the MES test in VPA-treated rats are shown in **Table 3**. Low-dose VPA offered moderate protection from seizures in rats within the first-hour post-dose using the MES test **(Figure 2C)**. High-dose VPA treatment in rats showed full to moderate seizure protection within two hours after drug treatment **(Table 3)**. In both dosing cohorts, protection was lost rapidly, consistent with the rapid elimination of VPA.

**Figure 2:**
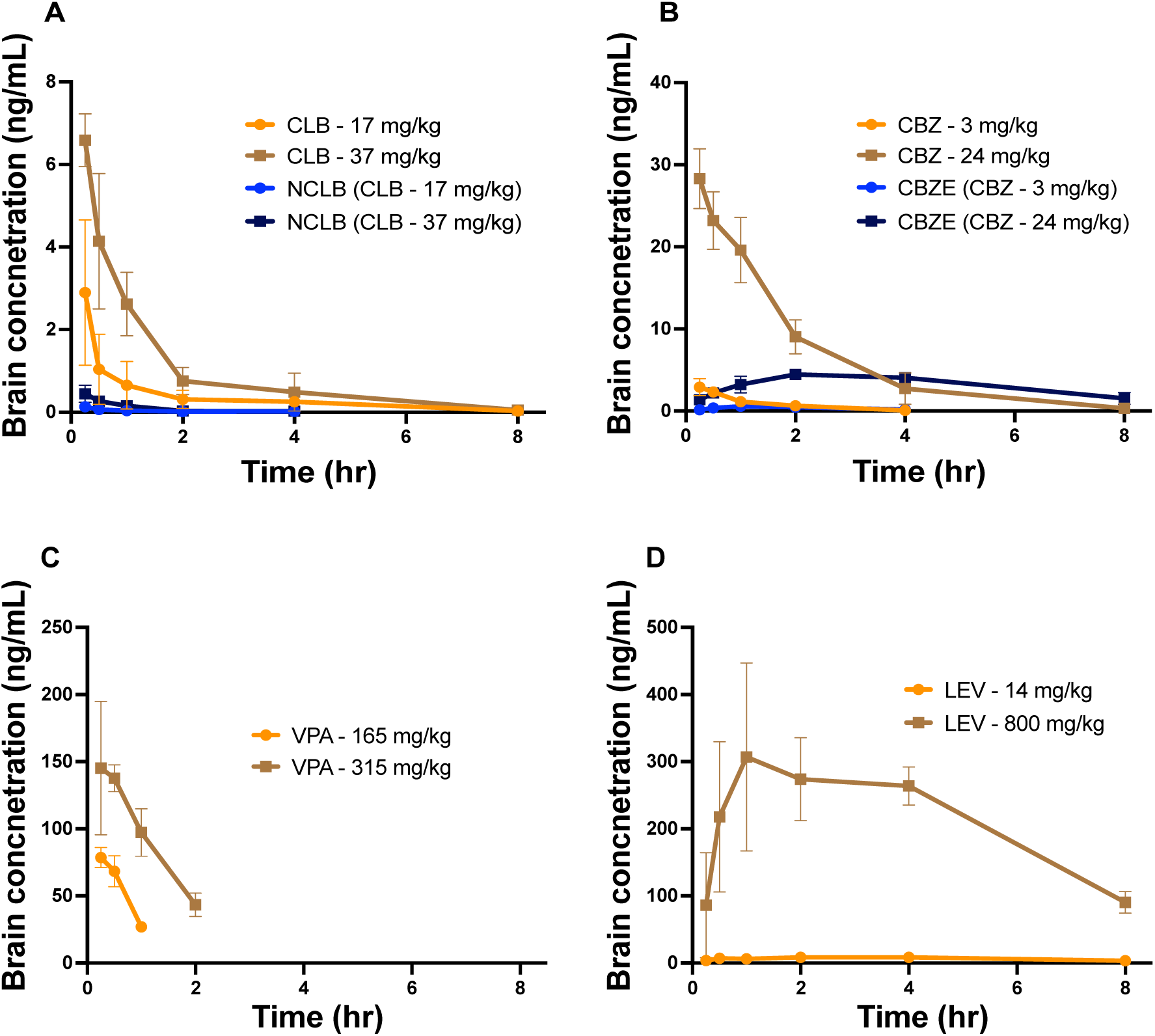
Concentration-time curves of (A) CLB and NCLB, (B) CBZ and CBZE, (C) VPA, and (D) LEV in the brain of adult male Sprague Dawley rats following intraperitoneal administration. Brain samples were obtained between 0-8 hr post-dose. Each data point is the mean brain concentration value for 4 rats.

### LEV

The PK parameters of LEV in both plasma and brain matrices are shown in **Table 4**. LEV t_1/2_ was approximately 2 hrs in mouse plasma and brain. LEV appears to be dose-proportional in mice, despite a 25-fold dose range. Seizure protection data from the 6 Hz 44 mA test in mice is depicted in **Table 2**, and the PK profile of LEV brain concentrations is shown in **Figure 1D**. The low-dose of LEV did not protect mice from seizures. The high-dose arm of LEV provided 25% protection against seizures within 4 hrs post-dose.

Plasma and brain t_1/2_ of LEV were longer in rats than in mice. In general, LEV exposures were higher in rats than in mice. **Figure 2D** shows brain levels of LEV in rats. Both low-and high-dose LEV treatments failed to protect against seizures at any time-point tested in the 6 Hz 80 V assay in rats **(Table 3)**

## Discussion

Epilepsy is a common neurologic condition linked with a substantial healthcare burden.^16,17^ Despite the advances in searching for more efficacious ASMs, about 30% of patients remain refractory to available therapies.^18^ Preclinical animal models can predict new compounds’ efficacy, safety, and initial PK profile before their introduction to humans^19^. Moreover, preclinical models that include PK evaluation can inform optimal dosing strategies and confirm brain exposure sufficient for target engagement. Therefore, the findings of this study support the use of preclinical PK evaluation combined with efficacy testing to provide a framework for subchronic dosing strategies, i.e., dosing over five days to 2 weeks for seizure protection.

Regarding chronic brain disorders such as epilepsy, their medical needs encompass a wide age-related range of patients and may affect various facets of drug design and dosing regimen.^20^ Effective treatment of seizures will most likely require well-designed long-term drug treatment. Hence, pre-clinical drug screening should include a subchronic dosing paradigm to mimic the clinic’s dosing regimen. Additionally, ASMs must cross the blood-brain barrier to reach their target cells. Therefore, it is essential to determine brain and plasma PK profiles of ASMs useful to inform appropriate doses for subchronic dosing.

Plasma and brain exposures of CLB and its clearance are similar between rats and mice.^21,22^ N-demethylation of CLB to NCLB is the main metabolic pathway. ^23,24,25^ The rate of disappearance of NCLB in humans, rats, and mice all differ which likely corresponds to the variations in CLB efficacy observed in these species.^21,26^ Our data indicate that higher NCLB concentrations were observed in mice than in rats due to differential metabolite formation and/or elimination.^21^ In mice, immediate post-dose efficacy of CLB was driven by the parent drug, but due to the rapid elimination of CLB, the persistence of beneficial effect was caused by NLCB due to the metabolite’s longer half-life. NCLB had been reported to contribute to the intensity and prolonged duration of CLB efficacy in rodents and humans.^26,27^ Data from our study suggest the high rate of NCLB formation in mice coupled with a longer half-life of NCLB in plasma and brain may contribute to the longer duration of action observed with CLB treatment in mice. In low-dose CLB treated rats, CLB failed to protect against seizures in the MES test. This observation is consistent with Hamilton et al.^25^ where CLB was only effective by the MES in rats at the median minimal toxic dose. CLB concentrations in rats are reported to be short-lived due to their rapid clearance.^26^ In addition, the overall lower concentrations of NCLB in plasma and brain of rats may have contributed to the lower efficacy of CLB in rats relative to mice.

Generally, the PK of CBZ in rodents is comparable to that of humans. Metabolic clearance via CYP2B6 and CYP3A4 are significant pathways for CBZ elimination, which is also impacted by autoinduction.^28,29,30^ The reported PK parameters in this study largely agree with previous findings. CBZ and CBZE generally displayed similar PK profiles between mice and rats. When comparing mice to rats, mice had a shorter CBZ half-life but a longer CBZE half-life than rats.^30,31^ The longer half-life of CBZE coupled with its high plasma and brain exposures in mice and rats may suggest that the metabolite contributes to the overall seizure protection benefits of CBZ in rats and mice.^32,33^ Given the longer half-life of CBZE and the known autoinduction of CYP2B6 and CYP3A4 by CBZ, we anticipate CBZE concentrations to accumulate during chronic dosing regimens.^34^ In rats, high-dose CBZ exposure was two-fold greater when dose adjusted in plasma and brain than in low-dose CBZ, suggesting a potentially saturable clearance mechanism at higher CBZ concentrations.^35^ We also observed a similar correlation between mice and rats in CBZ/CBZE seizure protection and PK profile. Additionally, our findings also suggest that the high CBZE exposure concentrations in plasma and brain, coupled with its longer half-life, may partially contribute to the total duration of action of CBZ.^36,37^

VPA demonstrates wide inter-species variability in its PK profile.^38,39^ VPA had a shorter half-life in rats than in mice. The differences in the half-life of VPA between species could be attributed to the differences in species-dependent drug-metabolizing enzymes.^40,41^ Our data are consistent with findings from Semmes and Shen ^43^, showing that plasma Cmax of VPA is dose-proportional in mice. We observed relatively low brain concentrationsof VPA in the high-dose group, which may be associated with capacity-limited brain sequestration in mice. ^42,43,44^ In low-dose VPA treatment in mice and rats, a decline in efficacy corresponded with rapid clearance of the drug in plasma and brain. Our data suggest that VPA had a short half-life in rats and mice and thus, indicates that VPA may require more frequent dosing in a chronic dosing regimen to achieve and maintain a steady-state therapeutic concentration range.

Our data indicate that half-lives of LEV in rats were relatively long in both plasma and brain. A longer half-life of LEV in the brain explains why LEV remains in the central nervous system (CNS) compartment for extended periods.^45,46^ LEV was dose-proportional, even across a 25-fold dose range, suggesting the absence of saturable clearance at physiologically relevant doses.^45^ We also observed a relatively high concentration of LEV in the brain of mice and rats. Our finding is supported by Doheny et al.,^47^ who found that LEV rapidly and readily crosses the blood-brain barrier to achieve comparable brain concentrations, which increase linearly and dose-dependently with serum concentration. LEV is ineffective in the MES and the 6 Hz 80 V 13 tests in mice and rats.^13,46^ In the present study, we tested LEV using high stimulus intensities, 44 mA and 80 V, of 6 Hz tests in mice and rats, respectively. However, we selected the ED_50_ of LEV from lower stimulus intensity 32 mA and 60 V for 6 Hz tests in mice and rats, respectively^5,13.^ Low-dose LEV treatment in mice tested in the 6 Hz 44 mA test failed to offer protection against seizure. Although our findings show detectable plasma and brain concentrations of LEV at a low dose, the achieved concentrations were likely below the therapeutic target ranges for seizure protection in the high stimulus seizure test in mice. We observed relatively prolonged but moderate seizure protection in mice at a high-dose LEV treatment. On the contrary, at the doses of LEV given in rats, LEV failed to offer protection against seizure at any time post-dose using the high stimulus of 80 V for 6 Hz, indicating that LEV is ineffective at the doses tested in the rat 6 Hz 80 V assay. Thus, the PK-guided preclinical studies on ASMs such as LEV are essential in designing dosing regimens and identifying appropriate models to evaluate efficacy to aid in proof-of-concept studies.

All data shown are from CF-1 mice and Sprague-Dawley rats, limiting this study’s ability to evaluate potential PK differences between mice and rat strains.^49^ However, these species represent two frequently used rodent species for testing ASMs. Additionally, there may be differences in pharmacological profiles of ASMs between various rodent seizure models; thus, the potency or antiseizure effect of some compounds at the selected doses may depend on the nature of the seizure induction assay and the subsequent mechanism(s) intrinsic to disease-induced brain modifications.^50^ Each time point represents pooled data from 4 animals in this study rather than collecting longitudinal samples from a single animal. This prevents an evaluation of inter-animal variability in PK. Brain tissue was not perfused with saline before homogenization, meaning residual blood in the sample may influence the observed brain concentration

The present study illustrates PK data’s utility in explaining the observed seizure protection time course and dose-response in rats and mice. Our findings indicate that species-specific variations in PK profiles may underlie the differences in observed efficacy of ASM in rats and mouse seizure models. Our data also demonstrate the importance of understanding the contribution of active metabolites to ASM efficacy. The study also sets the stage for exploring emerging subchronic dosing strategies.

## Acknowledgment

The authors thank the Epilepsy Therapy Screening Program (ETSP) at the National Institute of Neurological Disorders and Stroke (NINDS) for their review and comments on this manuscript.

## Authors Contribution

Study design and results interpretation: Jeffrey A. Mensah, Christopher A. Reilly, Karen S. Wilcox, Joseph E. Rower, Cameron S. Metcalf. Data analysis: Jeffrey A. Mensah, Joseph E. Rower, Cameron S. Metcalf. Data Collection: Jeffrey A. Mensah, Kristina Johnson, Joseph E. Rower.

## References

1. Chen Z, et al. Treatment Outcomes in Patients With Newly Diagnosed Epilepsy Treated With Established and New Antiepileptic Drugs: A 30-Year Longitudinal Cohort Study. JAMA Neurology, 2018. 75(3):279–286.

2. Wahab A. Difficulties in Treatment and Management of Epilepsy and Challenges in New Drug Development. Pharmaceuticals (Basel), 2010. 3(7):2090–2110.

3. Löscher W. Review: Critical review of current animal models of seizures and epilepsy used in the discovery and development of new antiepileptic drugs. Seizure, 2011. 20(5):359–368.

4. Rogawski MA. Review: Diverse mechanisms of antiepileptic drugs in the development pipeline. Epilepsy Research, 2006. 69(3):273–294.

5. Barton ME, et al. Pharmacological characterization of the 6 Hz psychomotor seizure model of partial epilepsy. Epilepsy Research, 2001. 47(3):217–227.

6. Rowley NM, White HS. Comparative anticonvulsant efficacy in the corneal kindled mouse model of partial epilepsy: correlation with other seizure and epilepsy models? Epilepsy Res, 2010. 92(2-3):163–169.

7. Markowitz GJ, et al. The pharmacokinetics of commonly used antiepileptic drugs in immature CD1 mice. Neuroreport, 2010. 21(6):452–456.

8. Esneault E, Peyon G, Castagné V. Efficacy of anticonvulsant substances in the 6 Hz seizure test: Comparison of two rodent species. Epilepsy Research, 2017. 134:9–15.

9. Bialer M, Twyman RE, White HS. Correlation Analysis between anticonvulsant ED_50_ values of AED drugs in mice and rats and their therapeutic doses and plasma levels. Epilepsy Behaviour, 2004. 5(6):866–872.

10. Kagan L, Zhao J, Mager DE. Interspecies Pharmacokinetic Modeling of Subcutaneous Absorption of Rituximab in Mice and Rats. Pharmaceutical Research, 2014. 31:3265–3273.

11. White HS, et al. General principles: discovery and preclinical development of antiepileptic drugs. 5th ed. Antiepileptic drugs, ed. R. H. Levy, et al. 2002, Philadelphia: Lippincott Williams & Wilkins.

12. Finney D. Probit analysis. A statistical treatment of the sigmoid response curve. 1952, Cambridge: University Press.

13. Metcalf C, et al. Development and pharmacologic characterization of the rat 6 Hz model of partial seizures. Epilepsia, 2017. 58(6):1073–1084.

14. Song M-X, et al. Synthesis and Evaluation of the Anticonvulsant Activities of 4-(2-(Alkylthio)benzo[d]oxazol-5-yl)-2,4-dihydro-3H-1,2,4-triazol-3-ones. Molecules, 2018. 23:756.

15. Isoherranen N, et al. Anticonvulsant profile of valrocemide (TV 1901): a new antiepileptic and CNS drug. Epilepsia, 2001. 42:831–836.

16. Devinsky O, et al., Epilepsy. Nature Reviews Disease Primers, 2018. 4(18024).

17. Thijs RD, et al, Epilepsy in adults. Lancet, 2019. 393:689–701.

18. Koneval Z, et al. Antiseizure drug efficacy and tolerability in established and novel drug discovery seizure models in outbred vs inbred mice. Epilepsia, 2020. 61(9):2022–2034.

19. Smith M, Wilcox KS, White HS. Discovery of Antiepileptic Drugs. Neurotherapeutics, 2007. 4(1):12–17.

20. Steinmetz K, Spack E. The basics of preclinical drug development for neurodegenerative disease indications. BMC Neuro, 2009. 9.

21. Caccia S, et al. Species differences in clobazam metabolism and antileptazol effect. J Pharm pharmacol, 1980. 32:101–103.

22. Sommerfeld-Klatta K, et al. New Methods Used in Pharmacokinetics and Therapeutic Monitoring of the First and Newer Generations of Antiepileptic Drugs (AEDs).. Molecules, 2020. 25(21):5083.

23. Tolbert D, Larsen F. Comprehensive Overview of the Clinical Pharmacokinetics of Clobazam. Clin Pharmacol., 2019. 59(1):7–19.

24. Giraud C, et al. In vitro characterization of clobazam metabolism by recombinant cytochrome P450 enzymes: importance of CYP2C19. Drug Metab Dispos, 2004. 32(11):1279–1286.

25. Hamilton K, et al. Persistent Hypersomnolence Following Clobazam in a Child With Epilepsy and Undiagnosed CYP2C19 Polymorphism. J Pediatr Pharmacol Ther., 2020. 25(4):320–327.

26. Volz M, et al. Kinetics and metabolism of clobazam in animals and man. Br. J. Clin. Pharmac, 1979. 7:41s–50s.

27. Haigh J. et al. N-desmethylclobazam: a possible alternative to clobazam in the treatment of refractory epilepsy? Br J Clin Pharmacol., 1987. 23(2):213–218.

28. Masuda Y, et al. Pharmacokinetic and pharmacodynamic tolerance of a new anticonvulsant agent (3-sulfamoylmethyl-1,2-benzisoxazole) compared to phenobarbital, diphenylhydantoin and carbamazepine in rats. Arch Int Pharmacodyn Ther., 1979. 240: 79–89.

29. Farghali-Hassan, et al. Carbamazepine pharmacokinetics in young, adult and pregnant rats. Relation to pharmacological effects. Arch Int Pharmacodyn Ther., 1976. 220:125–139.

30. Löscher W. The Pharmacokinetics of Antiepileptic Drugs in Rats: Consequences for Maintaining Effective Drug Levels during Prolonged Drug Administration in Rat Models of Epilepsy. Epilepsia, 2007. 48(7):1245–1258.

31. Remmel R, Sinz M, Cloyd J. Dose-dependent pharmacokinetics of carbamazepine in rats: determination of the formation clearance of carbamazepine-10,11-epoxide. Pharm Res, 1990. 7(5):513–517.

32. Kudriakova T, et al. Autoinduction and steady-state pharmacokinetics of carbamazepine and its major metabolites. Br. J. clin. Pharmac, 1992. 33:611–615.

33. Hönack D, Löscher W. Amygdala-kindling as a model for chronic .. efficacy studies on antiepileptic drugs: experiments with carbamazepine. Neuropharmacology, 1989. 28: 599–610.

34. Bertilsson L, Tomson T. Clinical pharmacokinetics and pharmacological effects of carbamazepine and carbamazepine-10,11-epoxide. An update. Clin Pharmacokinet, 1986. 11:177–198.

35. Tomson T, Tybring G, Bertilsson, L. Single-dose kinetics and metabolism of carbamazepine-10,11-epoxide. Clin. Pharmac. Ther., 1983. 33:58–65.

36. Chbili C, et al. The relationship between pharmacokinetic parameters of carbamazepine and therapeutic response in epileptic patients. Arch Med Sci., 2017. 13(2):353–360.

37. Potter J, Donnelly A. Carbamazepine-10,11-epoxide in therapeutic drug monitoring. Ther Drug Monit., 1998. 20(6):652–657.

38. Ghodke-Puranik Y, et al. Valproic acid pathway: pharmacokinetics and pharmacodynamics. Pharmacogenet Genomics., 2013. 23:236–241.

39. Nakashima H, et al. Determination of the Optimal Concentration of Valproic Acid in Patients with Epilepsy: A Population Pharmacokinetic-Pharmacodynamic Analysis. PLoS ONE. 10(10):e0141266.

40. Löscher W. Rapid Determination of Valproate Sodium in Serum by Gas-Liquid Chromatography. Epilepsia, 1977. 18(2):225–227.

41. Wilkinson G, Shand D. A physiological approach to hepatic drug clearance. Clinical Pharmacology & Theapeutics, 1975. 18(4):377–390.

42. Goldberg M, Todoroff T. Brain binding of anticonvulsants: carbamazepine and valproic acid. Neurology 1980. 30:826–831.

43. Semmes L, Shen D. Comparative Pharmacodynamics and Brain Distribution of E-A’-valproic and Valproate in Rats. Epilepsia, 1991. 32(2):232–241.

44. Hönack D, Löscher W. Intravenous valproate: onset and duration of anticonvulsant activity against a series of electroconvulsions in comparison with diazepam and phenytoin. Epilepsy Research, 1992. 13:215–221.

45. Patsalos P. The pharmacokinetic characteristics of levetiracetam. Methods Find Exp Clin Pharmacol, 2003. 25(2):123–129.

46. Steinhoff B, Staack A. Levetiracetam and brivaracetam: a review of evidence from clinical trials and clinical experience. Ther Adv Neurol Disord, 2019. 12.

47. Doheny H, et al. A comparison of the efficacy of carbamazepine and the novel anti-epileptic drug levetiracetam in the tetanus toxin model of focal complex partial epilepsy. Br J Pharmacol., 2002. 35(6):1425–1434.

48. Loscher W. Animal models of epilepsy for the development of antiepileptogenic and disease-modifying drugs. A comparison of the pharmacology of kindling and post-status epilepticus models of temporal lobe epilepsy. Epilepsy Res, 2002. 50(1-2):105–123.

49. Löscher W. The Search for New Screening Models of Pharmacoresistant Epilepsy: Is Induction of Acute Seizures in Epileptic Rodents a Suitable Approach? Neurochem Res 2017. 42:1926–1938.

50. Blanco, M., et al. Assessment of seizure susceptibility in pilocarpine epileptic and nonepileptic Wistar rats and of seizure reinduction with pentylenetetrazole and electroshock models. Epilepsia 2009. 50:824–831.

